# Repurposing a digital kitchen scale for neuroscience research: a complete hardware and software cookbook for PASTA

**DOI:** 10.1101/2020.04.10.035766

**Authors:** D Virag, J Homolak, I Kodvanj, A Babic Perhoc, A Knezovic, J Osmanovic Barilar, M Salkovic-Petrisic

**Author notes:** Corresponding author: Jan Homolak, MD.

## Abstract

Widely available low-cost electronics encourage the development of open-source tools for neuroscientific research. In recent years, many neuroscientists recognized the open science movement for its potential to stimulate and encourage science that is less focused on money, and more on robustness, validity, questioning and understanding. Here, we wanted to contribute to this global community by creating a research platform based on a common digital kitchen scale. This everyday ordinary kitchen tool is sometimes used in neuroscience research in various ways; however, its use is limited by sampling rate and inability to store and analyze data. To tackle this problem we developed a Platform for Auditory STArtle or PASTA. This robust and simple platform enables users to obtain data from kitchen scale load cells at a high sampling rate, store it and analyze it. Here, we used it to analyze acoustic startle and prepulse inhibition sensorimotor gating in rats treated intracerebroventricularly with streptozotocin, but the system can be easily modified and upgraded for other purposes. In accordance with open science principles, we shared complete hardware design with instructions. Furthermore, we also disclose our software codes written for PASTA data acquisition (C++, Arduino) and acoustic startle experimental protocol (Python) and analysis (R-based Awesome Toolbox for PASTA, ratPASTA R package). To further encourage the development of our PASTA platform we demonstrate its sensitivity by using PASTA-gathered data to extract breathing patterns during rat freezing behavior in our experimental protocol.

## Introduction

A recent substantial increase in the availability of low-cost electronics encouraged the development of open-source tools used in scientific research. The rapidly developing field of modern neuroscience is no exception, with a myriad of research groups openly publishing hardware and software with their peers with the goal of global advancement in the understanding of the neural basis of behavior. An agile acceptance of this open-science revolution stems from the core principles of open-science. First, in the context of the science reproducibility crisis, maximal transparency of the methodology, including bypassing “black box solutions” offered by manufacturers of equipment used in neuroscience experiments, should be considered as an important step forward. Moreover, projects released under open-source, *copyleft* licenses can, and should, be modified and improved by others, making the design more robust, scientifically verified, and easily adaptable for different experimental uses. Furthermore, open-source equipment is cost-effective, which makes it popular and widely used as it allows researchers to evade financial limitation-induced restraints and focus on science rather than money. Taken together, these concepts make open-source solutions indispensable for further evolution of neuroscience as they provide us with the tools to analyze what was previously unmeasured, and question what was taken as incontrovertible truth^1,2^. In this paper, we provide an open-source design for an experimental platform based on interfacing a microcontroller with a conventional digital kitchen scale. The utilization of a conventional digital kitchen scale for the collection of relevant scientific information has been used in the field of neuroscience for a long time, especially in the field of pain research and neurorehabilitation. For example, Matak elegantly used a digital kitchen scale to assess the magnitude of tetanus toxin-induced local rigidity of the rat gastrocnemius muscle^3^ and Koka and Hadlock reported that the kitchen scale-based extensor postural thrust test provides consistent quantitative results on functional recovery following rat sciatic nerve transection^4^. However, although conventional kitchen scales can provide valuable scientific information, their use is limited for several reasons. The experimenter using the kitchen scale has to carefully monitor the numeric output on the digital screen making the test prone to human error and bias. Moreover, although the sampling rate of the scale is relatively high (up to 80 Hz with our setup), human real-time information processing introduces a rate-limiting step making the output information rather modest both in regards to the amount of information and their temporal sequence. We argue that interfacing a kitchen scale with a computer would not only solve the aforementioned problems, but also introduce a whole new set of experimental opportunities such as rodent locomotor activity assessment or prepulse inhibition sensorimotor gating analysis. In the following text, we demonstrate this with our Platform for Acoustic STArtle - PASTA. Moreover, we describe the complete recipe with all hardware ingredients and ready-to-serve software packages, and test our system by a prepulse inhibition sensorimotor gating analysis in a rat model of sporadic Alzheimer’s disease.

## An overview of the Platform for Acoustic STArtle (PASTA)

The three key ingredients of PASTA are the aforementioned scale for registering the test animal’s activity, a loudspeaker to introduce the startling stimulus, and a computer acting as the control station which stores the scale data and automatically plays the startle stimulus through the loudspeaker when appropriate (Fig 1).

**Fig 1.**
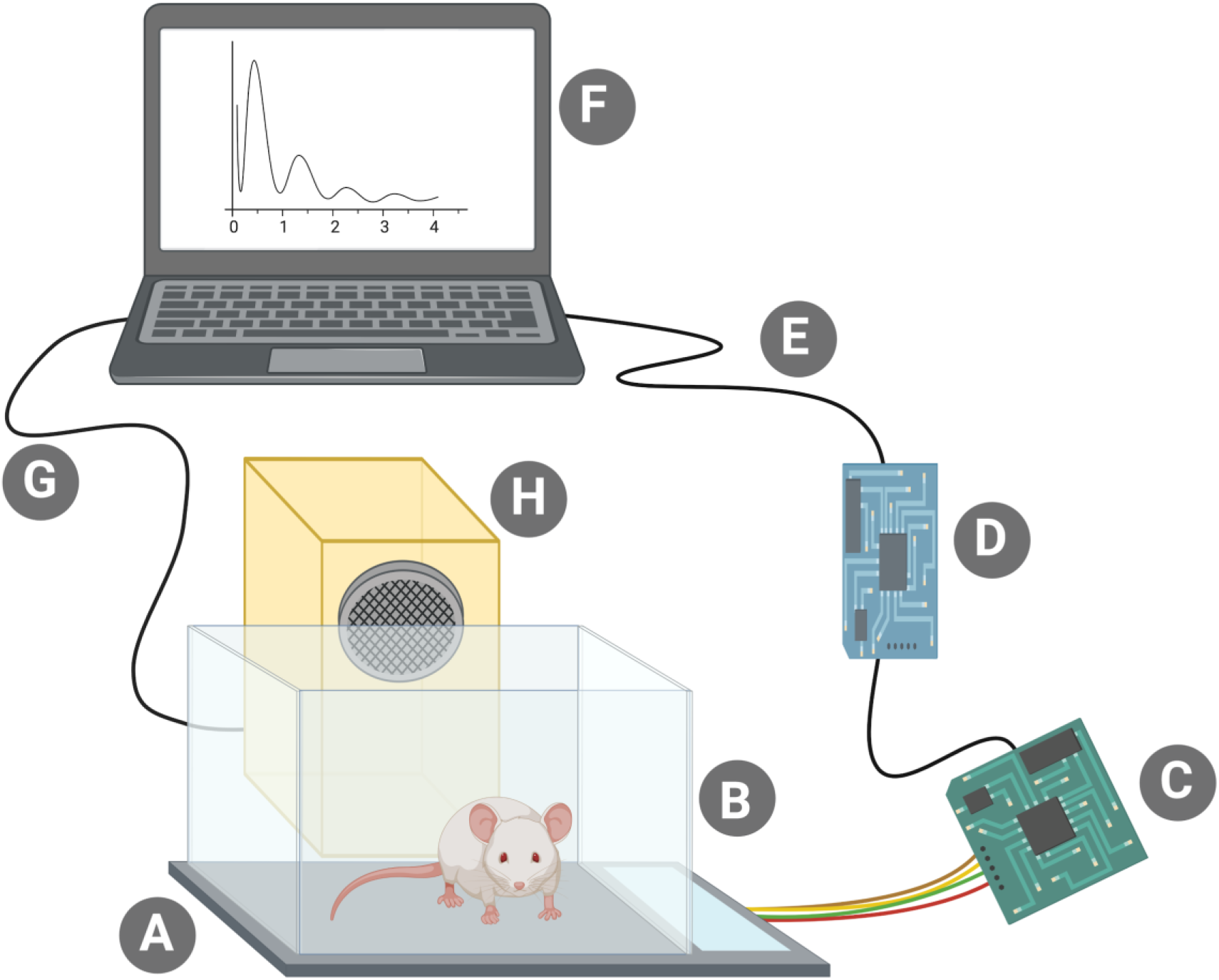
A schematic display of PASTA (Platform for Acoustic STArtle). **A)** The key ingredient of PASTA - a modified kitchen scale. **B)** A clean mouse cage affixed on top of the kitchen scale. **C)** A readily-available controller circuit board based on the HX711 integrated circuit **D)** A communication board, *NodeMCU ESP-32S* **E)** USB cable **F)** Your fancy laptop with our fancy Python script **G)** A normal audio cable. Really, there is nothing special about it. **H)** Standard loudspeaker.

**Fig 2.**
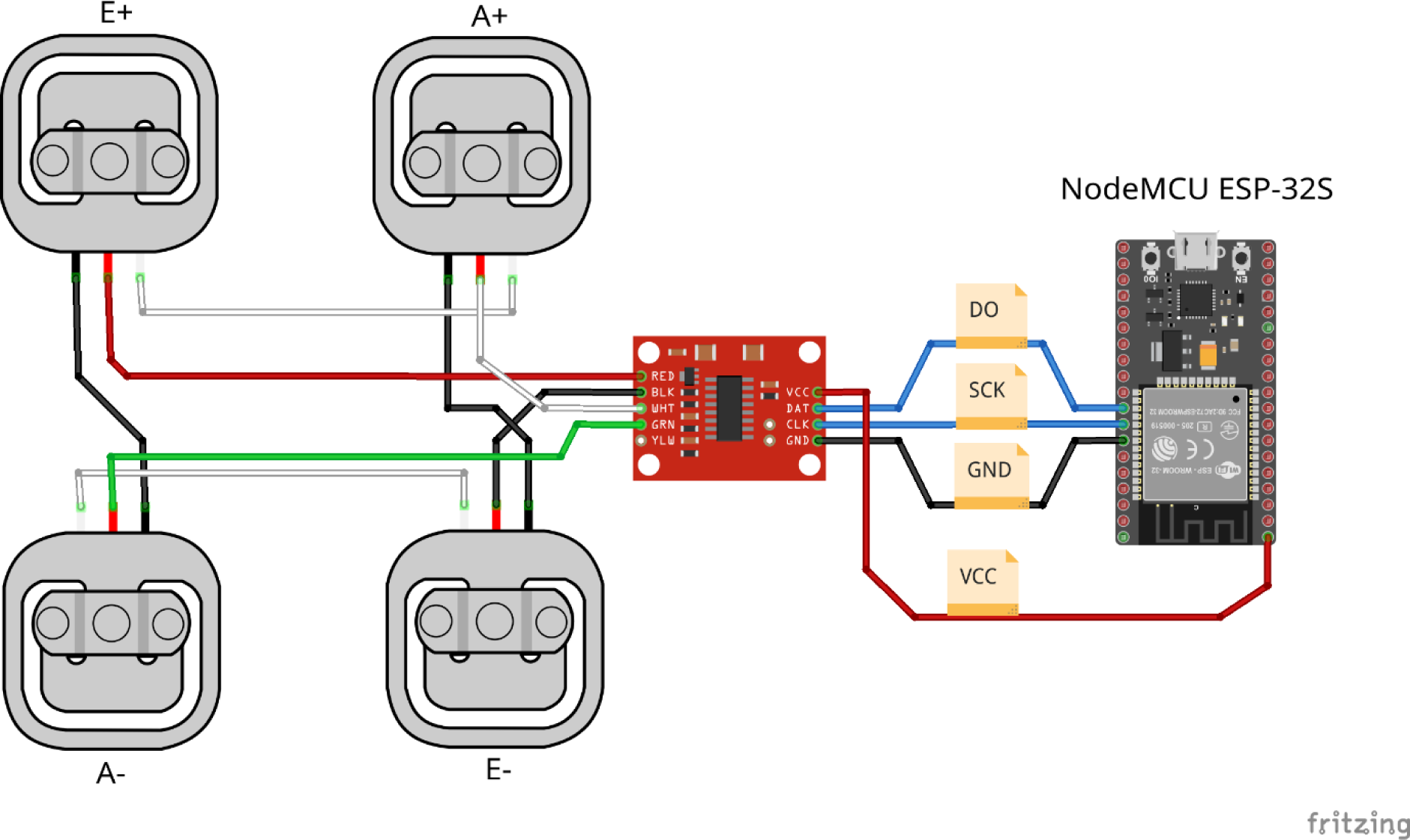
A schematic drawn in *Fritzing*^5^ representing how the four half-Wheatstone-bridge load cells found in our scale are wired together. They are then connected to their respective pins on the **HX711 circuit board**. This board communicates with the **NodeMCU ESP-32S** board through the DO and SCK lines routed to pins 18 and 19, respectively. The **NodeMCU ESP-32S** board provides power to the HX711 board and sends raw data to the computer over USB. This schematic is available in **Supplementary material**, as well as a full, annotated schematic made in KiCad.

## PASTA Hardware

To be able to read the Vivax Home KS-502T scale data at a high sampling rate and store it, we had to construct an appropriate interface between the load cells, which are the crux of every scale, and a computer. This interface uses a readily-available controller circuit board based on the HX711 integrated circuit (IC) to read the load cells’ resistance. The boards are usually configured for a 10 sample per second (SPS) sampling rate, so a hardware modification was applied which included reconnecting the HX711 *RATE* pin to *VCC* to enable the IC’s 80 SPS mode. The load cell controller board connects to a *NodeMCU ESP-32S* board which acts as a communication bridge and forwards the load cell data through its USB connection to the computer. As a bridge, more common AVR-based Arduino boards such as the Uno, Nano or Pro Micro may be used as well. The computer logs the received data and plays the startle sound at appropriate timestamps, which will be elaborated further in the *PASTA Software* section. As far as hardware considerations go, the choice of a speaker is the final point to note as the startling stimulus needs to be sufficiently loud to induce a startle response; however, the exact target volume depends heavily on the use case. The exact testing protocol we used in our test of this system can be found in **Supplement 1**.

### PASTA Software

Custom codes were written for the *NodeMCU ESP-32S* board (C++, Arduino), the control computer (Python), and for processing of collected data (R). All code can be found on corresponding GitHub repositories.

C++ code based on the Arduino framework which runs on the *NodeMCU ESP-32S* board simply receives samples from the HX711 IC and forwards it to the computer through its USB connection. Communication with the HX711 IC is done using the *HX711* library^6^, making the code portable to other targets specified in the library’s documentation, including typical AVR-based Arduino boards, simply by adjusting the appropriate pin definitions. The code can be found on GitHub^7^.

Before starting the experiment, a *WAVE* sound file named *startlesnd.wav* needs to be constructed for the entire duration of the experiment, with short pulses of white noise with or without prepulses, separated by a silence of appropriate duration, depending on the experiment that is being performed. We used *Audacity* to generate this file according to the specification detailed in **Supplement 1**. It is available as **Supplementary Audio Material 1**.

The computer runs Ubuntu, a Linux distribution, with PASTA Chef, a Python script on the computer, which automates the entire startle experiment and prepares *.pasta* files for further analysis. It starts by asking the user for the test animal’s name which will provide the data output file name, as well as the test animal’s mass, after which the experiment starts. Before starting, it is advisable to reset the communication bridge board using the on-board *RESET* button (labeled *EN* on the *NodeMCU ESP-32S board*). The sound file is loaded and playback is started automatically at the same time as data logging. During the experiment, scale data is also plotted on-screen in real time using *pyqtgraph*. After the experiment is done, all raw data files with the file extension *.pasta* can be found in the *data* directory, as well as a *mass.json* file which contains masses of all test animals. A caveat of this setup is that, without extensive kernel and sound system modification and reconfiguration, there is no way to precisely synchronize the recording to the timestamps in the data. This is further discussed in **Supplement 2**, and is a subject of our ongoing research. Startle latency can still be obtained by videotaping the experiment and correcting PASTA output according to the video. The code for PASTA Chef can be found on GitHub^8^.

A helper program, *C.A. V.A*^9^, was used to visualize the sound output on the computer screen, so that the exact moment when pulses are played can be registered during video analysis without relying on combined video-audio recordings.

To make data processing and visualisation easier, we have created an R package called rat PASTA (R-based Awesome Toolbox for PASTA), available on GitHub^10^. Briefly, the function *loadStartleData()*, loads and automatically merges all startle files from the folder. Unless otherwise specified, values are corrected for animal mass, and pulses are identified based on the inbuilt metadata. If necessary, users can manually specify which files to load and input custom metadata for puls identification. The functions *basicStartlePlot()* and *startlePlot()* are used to plot several graphs, whereas the function *summariseStartle()* returns a mathematical summary of the data. If users wish to expand the analysis, the output of the *loadStartleData()* function is a data frame and it can be used as an input for custom build functions. Detailed Information is provided on the website of the package^11^.

### A proof of concept study: prepulse inhibition sensorimotor gating analysis in a rat model of sporadic Alzheimer’s disease

In order to get a taste of PASTA, a prepulse inhibition sensorimotor gating analysis was conducted in rats previously treated intracerebroventricularly with streptozotocin (STZ-icv). This model has been extensively used to imitate the development of insulin resistant brain state in the sporadic form of Alzheimer’s disease^12^ and has been continually used by our group to study behavioral and neuropathological changes accompanying neurodegeneration. Prepulse inhibition (PPI) is an interesting cross-species neurobiological phenomenon describing the ability of the non-startling prepulse to weaken the subsequent startle response and has been mostly used to investigate sensorimotor gating in animal models of schizophrenia^13^. Furthermore, some researchers reported reduced PPI in transgenic animal models of Alzheimer’s disease^14^ and PPI deficits have been proposed as a potential biomarker of early Alzheimer’s disease^15^. On the other hand, others found no difference between patients and healthy controls^16^. To the best of our knowledge, PPI in STZ-icv rats was so far only examined by one group which found no difference between STZ-icv and control animals^17^. In the context of the reproducibility crisis we mentioned in the introduction, we seized this opportunity to both test PASTA and try to reproduce the findings originally reported by our colleagues from Brazil and Portugal. A complete detailed experimental protocol is available in **Supplement 2**. In short, twenty control (CTR) and twenty STZ-icv treated rats were placed in the PASTA system in a randomized order and the script described in the PASTA Software section was started. The script initiated PASTA data recording and activated the auditory sequence designed to provoke ten subsequent acoustic startle responses and ten prepulse-attenuated startles (**Supplementary Audio Material 1**). All gathered data was then processed by RAT PASTA. The whole procedure was filmed with a video camera, and a sample of the experimental procedure is available in **Supplementary Video Material 1**. Based on RAT PASTA analysis of the data obtained from PASTA, we conclude our platform and the auditory sequence we designed for this experiment were sufficient to register significant prepulse inhibition. Moreover, it is evident that both STZ-icv and CTR animals display significant PPI phenomenon with no apparent difference between the two groups (Table 1, Fig 3A), as already reported by Moreira-Silva et al.^17^. However, we also identified an intensified startle response to the initial pulses without prepulse signal in the STZ-icv group (Fig 3A). This is in accordance with behavioral abnormalities we previously observed in STZ-icv rats, and we plan to investigate the meaning of this interesting phenomenon further. Based on this proof of concept study, we conclude that the PASTA platform can be used to obtain meaningful neuroscientific information, and further development of our platform by us and other scientists involved with the open science community could further improve the platform and make it useful for a broad range of scientific experiments. Based on our preliminary testing, we already identified several additional pieces of information RAT PASTA and PASTA could provide with minimal adaptations.

One interesting example is based on the fact that our platform is sufficiently sensitive to register breathing patterns during freezing in rats (Fig 3B). A single breathing cycle registered by PASTA is depicted in Fig 3C-E. Corresponding thorax movements were calculated from the video file by baseline subtraction filtering, and frame pixel densitometry (Fig 3C-E). The whole analytical procedure is described in **Supplement 4** and baseline subtraction filtering is demonstrated in **Supplementary Video Material 2**. Representative breathing patterns can be observed in **Supplementary Video Material 3**. Raw data and data analysis protocols are available on GitHub^18^.

**Table 1.**
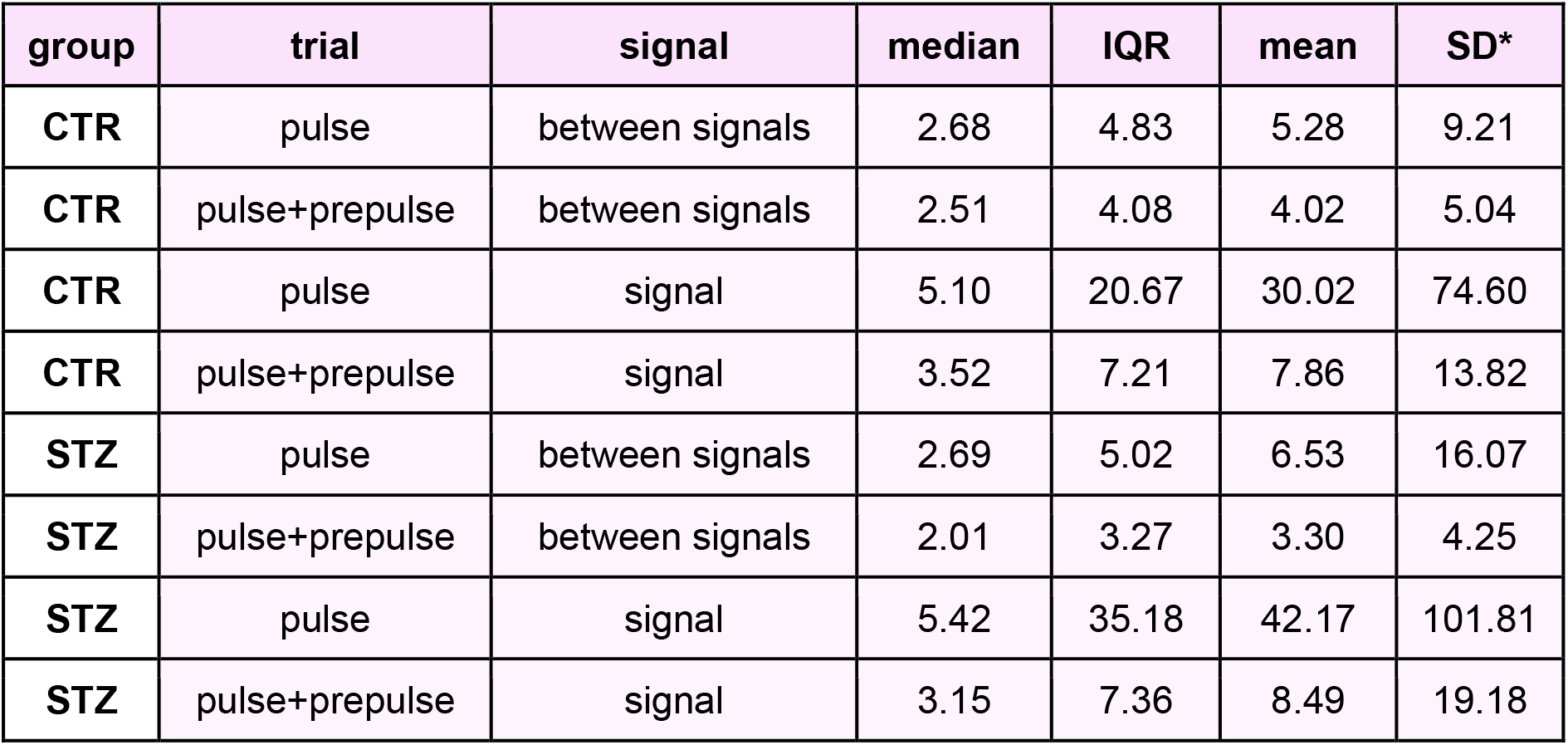
Descriptive statistics of the raw data obtained from PASTA. All results are presented in grams. *Very high standard deviation values of raw data obtained from PASTA indicate greater variation and in fact reflect the change we expected to see as the definition of the signal period is defined by the time window and includes both the signal and the noise recorded during the sampling segment. The best approach to quantification of PASTA-obtained changes is still to be determined, as a more detailed mathematical approach might increase the sensitivity and specificity of PASTA.

**Fig 3.**
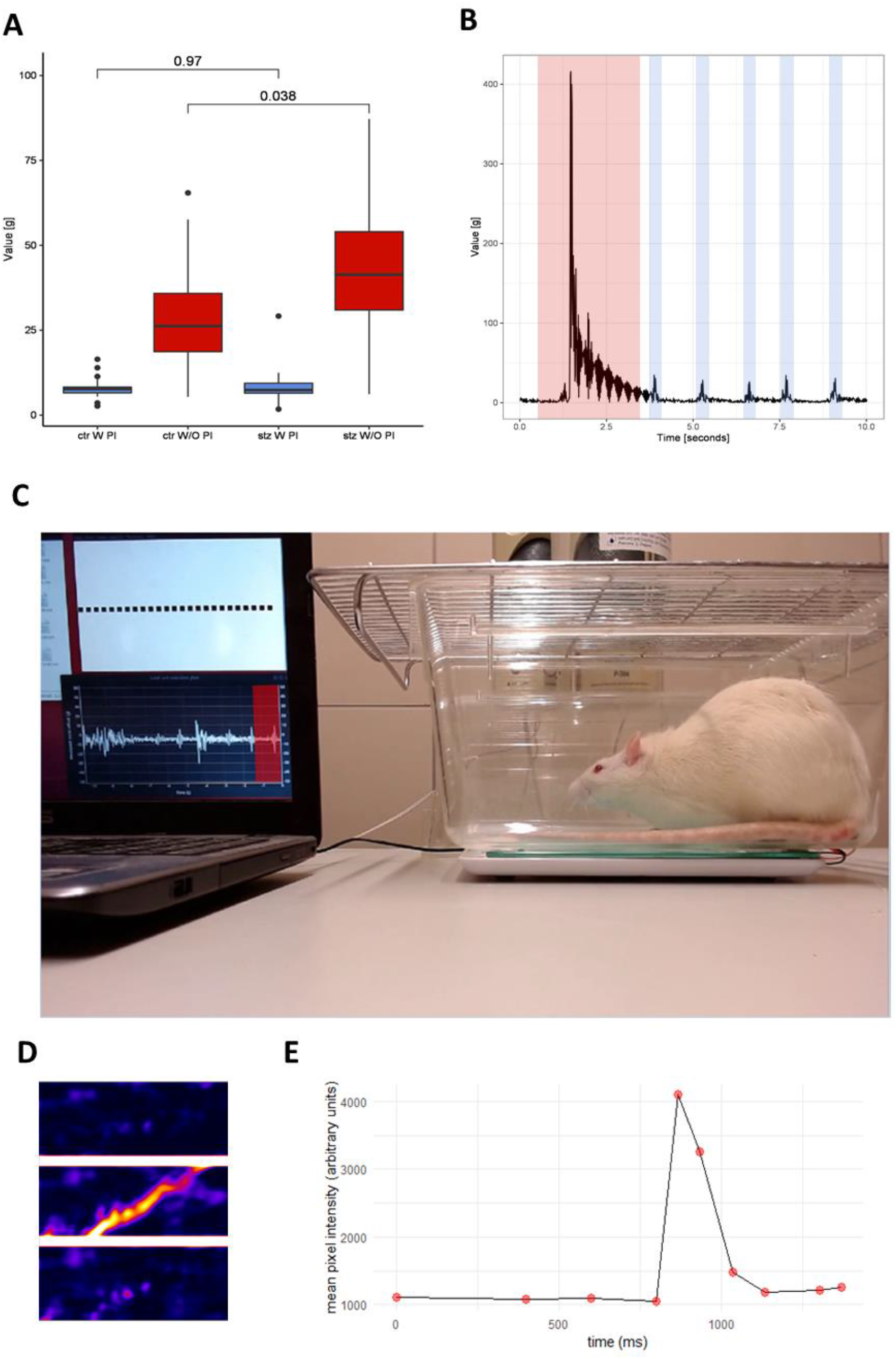
Results of the startle response and prepulse inhibition sensorimotor gating analysis obtained by the Platform for Acoustic STArtle (PASTA), PASTA Chef and R-based Awesome Toolbox for PASTA (ratPASTA). **A**) A box plot depiction of the startle response intensities for control (CTR) and rats treated with intracerebroventricular streptozotocin (STZ). W PI (With Prepulse Inhibition) indicates the response was calculated by pooling data obtained from the trials that included prepulse inhibitory stimulus, and W/O PI (Without Prepulse Inhibition) indicates the response was calculated by pooling data obtained from the trials that did not include prepulse inhibitory stimulus. Both CTR and STZ rats display the prepulse inhibition phenomenon, and STZ rats differ from the controls by the pronounced startle response to the initial stimulus not preceded by the inhibitory prepulse (p=0.038). **B**) Raw data display of a single startle response trial. The area in red indicates a startle response, and areas in blue indicate breathing patterns evident during the freezing behavior. **C**) Sample image from one trial during the proof of concept study. The laptop running the *PASTA Chef* script is displayed on the left side of the picture with C.A.V.A. helper program on the upper side of the screen and load cell real-time plot on the lower side of the screen. The red area indicates a single breathing pattern further examined by corresponding video analysis and depicted in Fig 3D-E. The green area indicates the thoracic area of interest used for further video and image analysis depicted in Fig 3D-E. **D**) Portrayal of a single breathing cycle obtained by baseline subtraction filtering video analysis. Sample images of a single cycle are arranged in appropriate temporal order with the uppermost image representing the baseline state before thoracic movement, and lowermost image the end of the movement. **E**) A visual representation of a breathing cycle obtained by densitometric analysis of the images obtained by baseline subtraction filtering video analysis.

**Fig 4.**
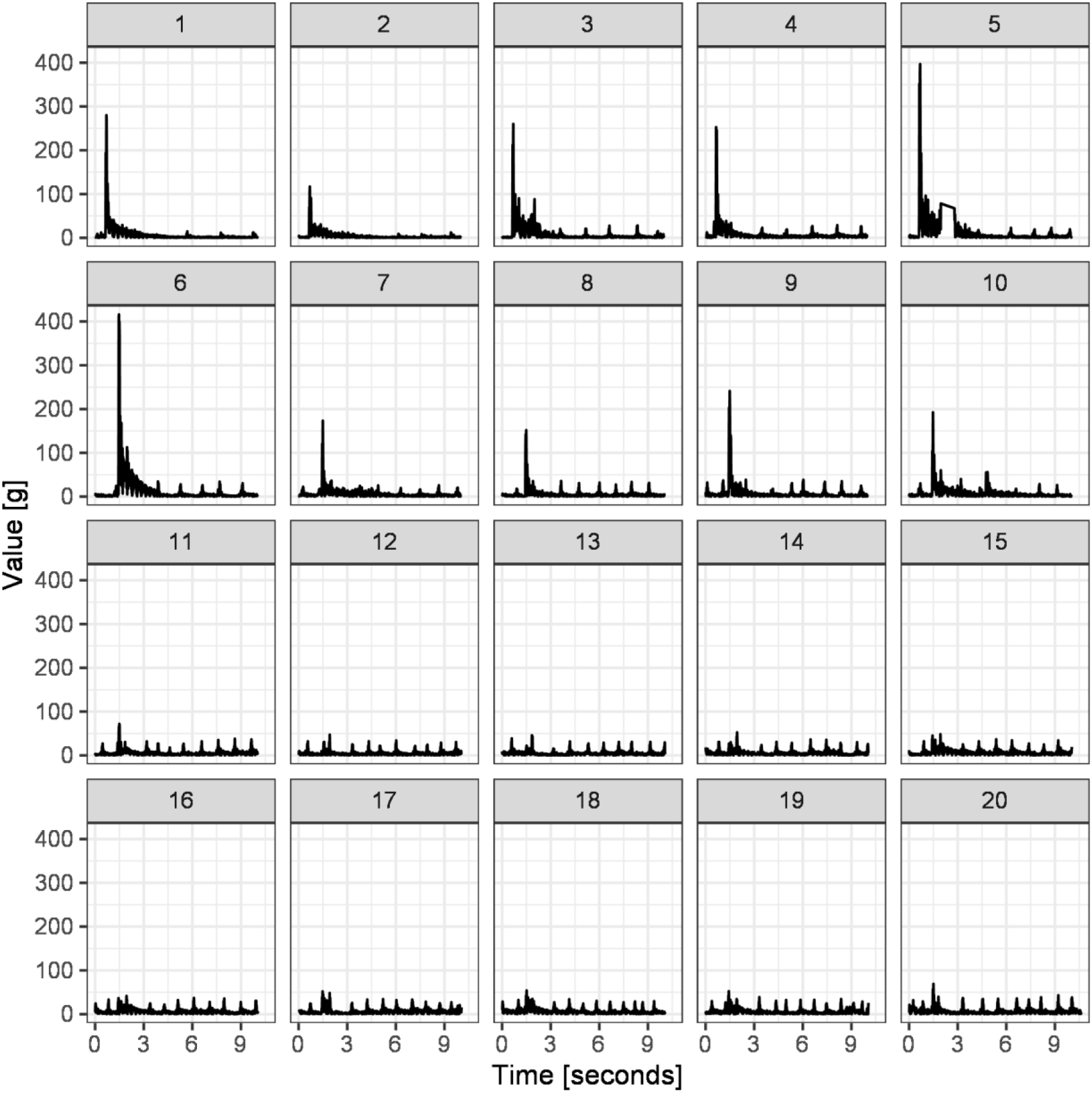
A representative example of the results obtained by the experimental protocol used for the acoustic startle and prepulse inhibition sensorimotor gating analysis. The experimental protocol and audio sequence are further discussed in the PASTA Software section and the acoustic startle sequence is available in **Supplementary Audio Material 1**. In short, trials 1-10 are focused on the quantification of the acoustic startle response, and trials 11-20 consist of the same startle stimulus preceded by the prepulse inhibitory stimulus, and are focused on the quantification of the prepulse inhibition phenomenon. Small peaks represent breathing patterns recorded during freezing behavior of the rat and are further discussed in Fig 3 with startle response indicated in red and breathing patterns indicated in blue in Fig 3B.

## Conclusion

In this brief paper, we introduce PASTA, a low-cost open-source platform for control of common kitchen scale load cells, which can be used to extract meaningful and important behavioral data. In accordance with principles of open science, we provide complete instructions on how to demolish your regular kitchen scale and use the parts to turn it into a beautiful multifunctional neurobehavioral platform. Moreover, we provide the whole code needed to design PASTA software - a C++ and Arduino code for the communication board, a Python code for the control computer, and even ratPASTA, an R-based package for data processing, analysis and visualization - so you can slightly modify everything to better fit your experiment, sit back and enjoy PASTA. In our proof of concept study, we have shown how PASTA can be used for acoustic startle and prepulse inhibition analysis, but the whole platform is easily adaptable, and we hope to see our colleagues using our platform for other types of behavioral analyses further building on PASTA in the spirit of open-science.

## Supporting information

Supplementary Fritzing Schematic

Supplementary KiCad Schematic

## Author’s contributions

**DV**, **JH** and **IK** conceptualized PASTA, contributed equally and leading roles in different parts of the project were equally distributed. **DV** is responsible for PASTA hardware design, kitchen scale hacking and writing the *NodeMCU ESP-32S* C++/Arduino software code and Python script for the acoustic startle experiment. **DV** and **JH** are responsible for the design and **JH** is responsible for the execution of the experimental protocol. **IK** is the main author responsible for the code behind the ratPASTA R-package and visualization used in the manuscript. **JH** is responsible for the video analysis. **DV**, **JH** and **IK** designed the analyses and wrote the manuscript. **JH**, **ABP**, **AK** and **JOB** generated a rat model of sporadic Alzheimer’s disease by intracerebroventricular administration of streptozotocin (STZ-icv). **MSP** is head of the Laboratory for Molecular Neuropharmacology where all experiments were conducted, principal investigator of the Croatian Science Foundation funded project (IP-2018-01-8938) and supervisor and mentor of **DV**, **JH** and **IK**. All authors approved the manuscript and provided comments during the writing process.

## Ethics committee approval

All experiments were conducted in concordance with the highest standard of animal welfare. Only certified personnel handled animals. Animal procedures were carried out at the University of Zagreb Medical School (Zagreb, Croatia) and were in compliance with current institutional, national (The Animal Protection Act, NN135/2006; NN 47/2011), and international (Directive 2010/63/EU) guidelines governing the use of experimental animals. The experiments were approved by the national regulatory body responsible for issuing ethical approvals, the Croatian Ministry of Agriculture and by the Ethical Committee of the University of Zagreb School of Medicine.

## References

1. Krakauer, J. W., Ghazanfar, A. A., Gomez-Marin, A., MacIver, M. A. & Poeppel, D. Neuroscience Needs Behavior: Correcting a Reductionist Bias. Neuron 93, 480–490 (2017).

2. White, S. R., Amarante, L. M., Kravitz, A. V. & Laubach, M. The Future Is Open: Open-Source Tools for Behavioral Neuroscience Research. eNeuro 6, (2019).

3. Matak, I. Evidence for central antispastic effect of botulinum toxin type A. Br. J. Pharmacol. 177, 65–76 (2020).

4. Koka, R. & Hadlock, T. A. Quantification of functional recovery following rat sciatic nerve transection. Exp. Neurol. 168, 192–195 (2001).

5. Fritzing. https://fritzing.org.

6. bogde. HX711. GitHub https://github.com/bogde/HX711.

7. davorvr. PASTA Bridge. GitHub https://github.com/davorvr/pasta-bridge.

8. davorvr. PASTA Chef. GitHub https://github.com/davorvr/pasta-chef.

9. karl. cava. GitHub https://github.com/karlstav/cava.

10. Kodvanj I., Virag D., Homolak J. ratPASTA. R package version 0.1.0. https://github.com/ikodvanj/ratPASTA (2020).

11. Website. Available at: https://ikodvanj.github.io/ratPASTAsite/index.html

12. Salkovic-Petrisic, M., Knezovic, A., Hoyer, S. & Riederer, P. What have we learned from the streptozotocin-induced animal model of sporadic Alzheimer’s disease, about the therapeutic strategies in Alzheimer’s research. J. Neural Transm. 120, 233–252 (2013).

13. Powell, S. B., Zhou, X. & Geyer, M. A. Prepulse inhibition and genetic mouse models of schizophrenia. Behav. Brain Res. 204, 282–294 (2009).

14. Sichler, M. E., Löw, M. J., Schleicher, E. M., Bayer, T. A. & Bouter, Y. Reduced Acoustic Startle Response and Prepulse Inhibition in the Tg4-42 Model of Alzheimer’s Disease. J Alzheimers Dis Rep 3, 269–278 (2019).

15. Ueki, A., Goto, K., Sato, N., Iso, H. & Morita, Y. Prepulse inhibition of acoustic startle response in mild cognitive impairment and mild dementia of Alzheimer type. Psychiatry Clin. Neurosci. 60, 55–62 (2006).

16. Hejl, A.-M., Glenthøj, B., Mackeprang, T., Hemmingsen, R. & Waldemar, G. Prepulse inhibition in patients with Alzheimer’s disease. Neurobiol. Aging 25, 1045–1050 (2004).

17. Moreira-Silva, D. et al. Anandamide Effects in a Streptozotocin-Induced Alzheimer’s Disease-Like Sporadic Dementia in Rats. Front. Neurosci. 12, 653 (2018).

18. davorvr. pasta-data. GitHub https://github.com/davorvr/pasta-data.

