## Supplement 1 for "Repurposing a digital kitchen scale for neuroscience research: a complete hardware and software cookbook for PASTA"

### Supplementary material 1

#### Volume considerations

To produce a startle response with a sound pulse, as well as startle inhibition with a prepulse, appropriate volumes need to be used which will elicit the desired response in the test animal. Valsamis, B & Schmid, S have found that rodents typically begin to startle at 85-90 dB, with the response peaking at 100-110 dB, and they recommend a prepulse volume of 75 or 85 dB[1]. As our proof of concept study was successful in eliciting startle and startle inhibition responses in rats, we obtained measurements from a similar setup, which is based on common, amplified desktop loudspeakers. Lacking expensive, calibrated professional audio equipment necessary to obtain a precise measurement, we used the open-source Android application *OpeNoise*[2] to reveal a volume of 88.9 dB for pulses, and 77.9 dB for prepulses. Users are advised to adapt their setup until obtaining a satisfactory response in test animals, in accordance with their specific use case.

#### Pulse/prepulse timing considerations

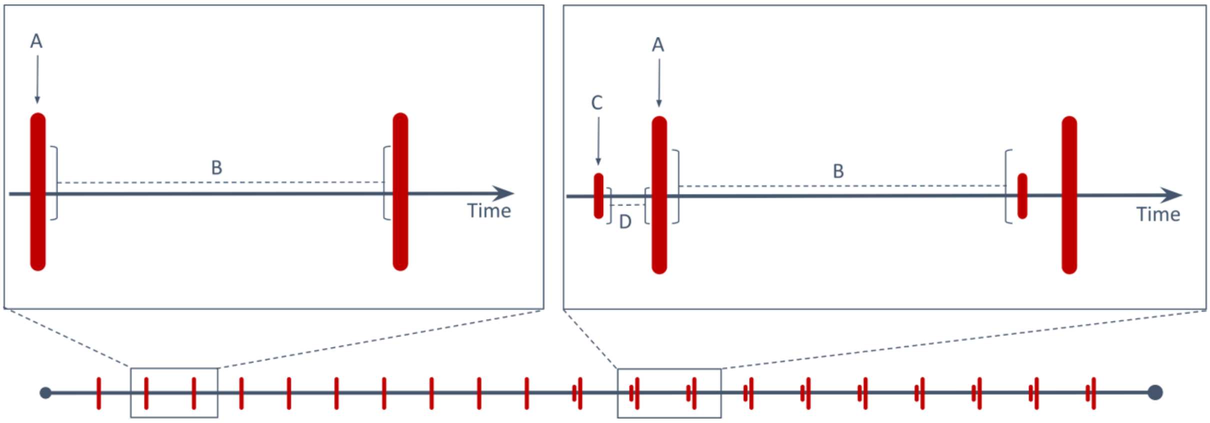


**Fig. 1** An auditory startle pulse consists of **A)** 20 milliseconds of white noise at 95% amplitude, followed by **B)** 10 seconds of silence. A prepulse occurs immediately before a pulse, and it consists of **C)** 4 milliseconds of white noise at 10% amplitude, followed by **D)** 30 milliseconds of silence.

As stated in the main text, before starting the experiment, a *WAVE* sound file named *startlesnd.wav* needs to be constructed with the appropriate number, duration and timing of pulses and prepulses. The protocol used in our proof of concept study consists of a starting 10 seconds of silence, followed by 10 pulses, 10 pulses with prepulses, and a 10-second silence at the end, with a total run time of 3 minutes and 30 seconds per animal. Detailed timings and amplitudes are shown in Fig 3. We used Audacity to generate this audio file, which is available as **Supplementary Audio Material 1**. Users are encouraged to modify and adapt this protocol according to the needs of their specific study.

1. Valsamis, B. & Schmid, S. Habituation and prepulse inhibition of acoustic startle in rodents. *J. Vis. Exp.* e3446 (2011).

2. OpeNoise meter. Github repository at https://github.com/Arpapiemonte/openoise-meter (5th April 2020).
