## Supplement 2 for "Repurposing a digital kitchen scale for neuroscience research: a complete hardware and software cookbook for PASTA"

### Supplementary material 2

As mentioned in the text, a notable, but correctable, limitation of this setup is a varying and unknowable latency that occurs when starting the audio using *aplay*. This delay is likely related to the interaction between software and hardware through the sound server PulseAudio and the Linux audio subsystem, ALSA. There are some mitigations that could be applied, such as running *PASTA Chef* on a Linux system with PREEMPT_RT patches[1] and without the PulseAudio audio server, but a solution likely lies in delegating the audio output, as well as scale reading, to an embedded system. A microcontroller running compiled C/C++ code without an underlying operating system or one with a real-time operating system such as FreeRTOS could adequately synchronize audio and data output. This kind of system would ensure a predictably timed audio output at predefined timestamps in *.pasta* data even without additional correction with video recordings, and is a subject of our ongoing and future research.

1. The Real Time Linux collaborative project. https://wiki.linuxfoundation.org/realtime/start.
