## Supplement 3 for "Repurposing a digital kitchen scale for neuroscience research: a complete hardware and software cookbook for PASTA"

### Supplementary material 3

Forty male Wistar rats, bred and kept in-house (Department of Pharmacology, University of Zagreb School of Medicine) were used for the experiment. The animals were kept in an animal facility with stable conditions (constant temperature and humidity – 22-24ºC and 40-60%, and a 12-h light/12-h dark cycle (7AM/7PM)) in standard cages with wood-chip bedding and food and water *ad libitum*; with three animals placed per cage. At the age of three months, the animals underwent intracerebroventricular treatment with either streptozotocin (STZ-icv; 20 animals) or 0.05M citrate buffer, pH 4.5 (CTR; 20 animals). STZ is a beta-cytotoxic substance selective for insulin-secreting/producing cells, which has been abundantly used to induce insulin-resistant brain state in laboratory animals, mimicking the main hallmarks of sporadic Alzheimer's disease. STZ was applied intracerebroventricularly to rats in deep anesthesia (ketamine 50 mg/kg /xylazine 5 mg/kg, ip) in a dose of 3 mg/kg divided in two doses (48 hours apart) into the rats' lateral ventricles (according to the procedure first described by Noble et al.[1] and used by our research team in previous experiments[2–4]), whereas control animals received only the vehicle, citric buffer, in the same manner. The proof of concept study for assessing the Platform for Acoustic STArtle (PASTA) was conducted 1 month after STZ-icv treatment. All animals were placed in the clean mouse cage affixed on top of the kitchen scale transformed into PASTA and closed from the above by a standard rat cage flooring grid one by one in a randomized order. Once the animal placed in the mouse cage stopped moving, the microcontroller was reset to calibrate load cell readings and the Python script *PASTA Chef* described in the main text was started. After the audio sequence finished, the animal was returned to its home-cage and the mouse cage affixed to the platform was cleaned with 70% ethanol before beginning the next trial. The whole procedure was performed in an isolated laboratory in order to assure only one animal at the time was exposed to the acoustic stimulus. All measurements were conducted between 11 AM and 3 PM. Animals were monitored during the next 24 hour to assess whether they were stressed out by the procedure. The animals were afterwards used for acute intracerebroventricular administration of the gastric inhibitory polypeptide receptor inhibitor Pro(3)GIP as part of the Croatian Science Foundation funded project "Mechanisms of nutrient-mediated effects of endogenous glucagon-like peptide-1 on cognitive and metabolic alterations in experimental models of neurodegenerative disorders (ID: IP-2018-01-8938; PI: Melita Šalković-Petrišić MD, PhD)" in accordance to 3R principles of animal welfare and following all ethical standards.

1. Noble, E. P., Wurtman, R. J. & Axelrod, J. A simple and rapid method for injecting H3-norepinephrine into the lateral ventricle of the rat brain. *Life Sci.* **6**, 281–291 (1967).
2. Knezovic, A. *et al.* Staging of cognitive deficits and neuropathological and

ultrastructural changes in streptozotocin-induced rat model of Alzheimer's disease. *Journal of Neural Transmission* vol. 122 577–592 (2015).

3. Grünblatt, E., Salkovic-Petrisic, M., Osmanovic, J., Riederer, P. & Hoyer, S. Brain insulin system dysfunction in streptozotocin intracerebroventricularly treated rats generates hyperphosphorylated tau protein. *J. Neurochem.* **101**, 757–770 (2007).
4. Knezovic, A. *et al.* Rat brain glucose transporter-2, insulin receptor and glial expression are acute targets of intracerebroventricular streptozotocin: risk factors for sporadic Alzheimer's disease? *J. Neural Transm.* **124**, 695–708 (2017).
