## Supplement 4 for "Repurposing a digital kitchen scale for neuroscience research: a complete hardware and software cookbook for PASTA"

### Supplementary material 4

As described in the main text, we recognized that the Platform for Acoustic STArtle (PASTA) was sufficiently sensitive to recognize breathing patterns during freezing behaviour in rats. In order to test whether signals obtained from PASTA really matched breathing patterns, we used video recordings of PASTA proof of concept study trials. All trials from the proof of concept experiment were recorded by Logitech C920 HD PRO webcam and Webcamoid software available on GitHub[1]. A single video file from the experiment was used (STZ-icv animal number 16) to examine how thorax movements from a random part of the video trial correspond to PASTA-recorded breathing signals. The video file was first processed by HandBrake[2] software in order to force a constant frame rate (30 frames per second) and ensure better temporal accuracy. A high quality video file was exported in MP4 format and processed with Tracker Video Analysis and Modeling Tool[3]. In short, the video was set to begin during the rat's freezing behavior, a pointer was used to apply maximal zoom over the thoracic area and baseline filtering was enabled to subtract baseline image and display only pixels with changing intensities (**Supplementary Video Material 2**). The video was then manually played frame by frame with inter-frame durations of 33,33 ms. The number of frames necessary to describe one cycle of inspiration-expiration was assessed to be approximately 10 measurements per cycle of approximately 1,5 s. Frames of interest were exported as image files and further processed by Fiji software[4] by using the built-in "Measure" function and mean pixel intensity values were used for further visualization in R by means of the ggplot2 package[5,6]. PASTA-recorded raw values of the corresponding breath were marked in red on the screen of the computer running the Python script *PASTA Chef* and the thoracic region of interest was marked in green (Fig 3C).


1. webcamoid. webcamoid/webcamoid. *GitHub* https://github.com/webcamoid/webcamoid.

2. HandBrake: Open Source Video Transcoder. https://handbrake.fr/.

3. Tracker Video Analysis and Modeling Tool for Physics Education.

https://physlets.org/tracker/.

4. Fiji. *ImageJ* https://imagej.net/Fiji.

5. R: The R Project for Statistical Computing. https://www.r-project.org/.

6. Create Elegant Data Visualisations Using the Grammar of Graphics [R package ggplot2 version 3.3.0].
