## Supplementary KiCad Schematic for "Repurposing a digital kitchen scale for neuroscience research: a complete hardware and software cookbook for PASTA": pasta-scale.pdf

### Title: PASTA Scale – full, annotated schematic

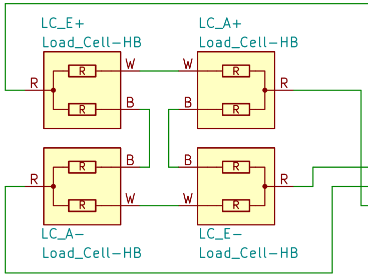


This is a schematic which shows how the load cells need to be connected. The symbols are positioned to have the wiring between the cells visualized as cleanly as possible. Be sure to keep precise track of the colors! Note that the A+ line only crosses E– and A– lines. It is NOT connected to them!

Here is a hopefully helpful diagram:

BLACK connects opposite letters AND signs:
E+ <––BLACK––> A–
E– <––BLACK––> A+

WHITE connects opposite letters to same signs:
E+ <––WHITE––> A+
E– <––WHITE––> A–

RED connects to the cells' respective connections on the HX711 module.

These components usually come as part of an HX711–based module. This layout is based on SparkFun's Load Cell Amplifier (SEN–13879).

NOTE: The RATE pin is connected to GND on most HX711–based modules. To enable the IC's 80 SPS mode, RATE needs to be connected to VCC, as shown here.

NOTE: The authors are not affiliated with SparkFun Electronics in any way, shape or form.

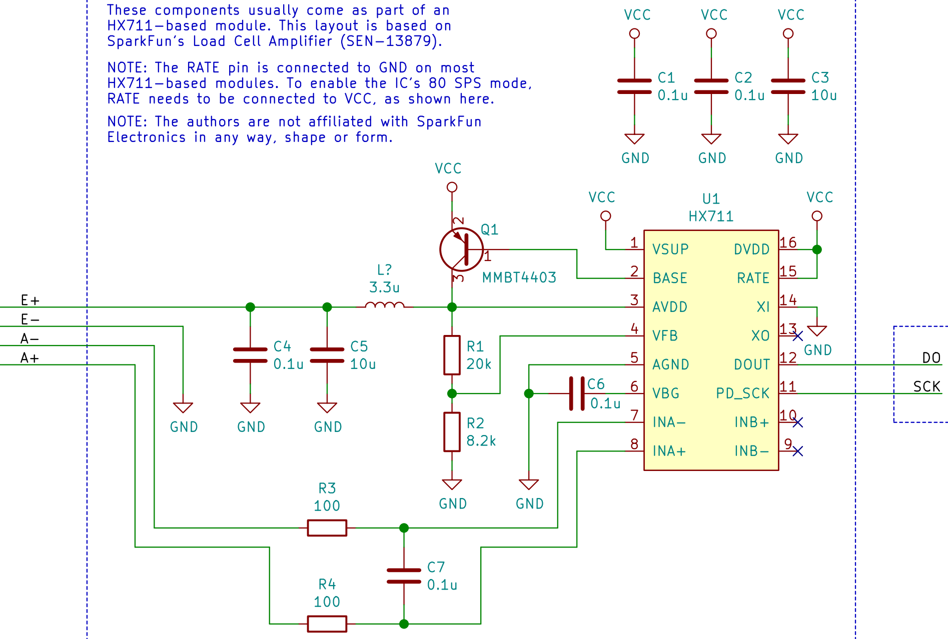


These lines connect to PASTA Bridge, which is simply an Arduino–based microcontroller board that forwards HX711 data to a PC over a USB serial interface.

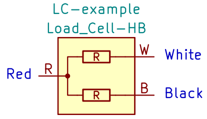


This part is a half–bridge load cell, as found in our Vivax Home KS–502T kitchen scale. The wires are typically color–coded as shown, and the symbol contains this type of load cell's equivalent circuit.

Schematic for all custom electronics used in the paper "Repurposing a digital kitchen scale for neuroscience research: a complete hardware and software cookbook for PASTA" by Virag D, Homolak J, Kodvanj I, et al.

DOI: https://doi.org/10.1101/2020.04.10.035766

**Department of Pharmacology, University of Zagreb School of Medicine, Zagreb, Croatia**

Sheet: /
File: pasta–scale.sch

***Title: PASTA Scale – full, annotated schematic***

| | | |
|---|---|---|
| Size: A4 | Date: 2020–04–16 | **Rev: v1** |
| KiCad E.D.A. kicad 5.1.5 | | Id: 1/1 |
