## Supplementary KiCad Schematic for "Repurposing a digital kitchen scale for neuroscience research: a complete hardware and software cookbook for PASTA": pasta-scale.pro

```
update=četvrtak, 16. travnja 2020. 19:07:12 CEST
version=1
last_client=kicad
[general]
version=1
RootSch=
BoardNm=
[pcbnew]
version=1
LastNetListRead=
UseCmpFile=1
PadDrill=0.600000000000
PadDrillOvalY=0.600000000000
PadSizeH=1.500000000000
PadSizeV=1.500000000000
PcbTextSizeV=1.500000000000
PcbTextSizeH=1.500000000000
PcbTextThickness=0.300000000000
ModuleTextSizeV=1.000000000000
ModuleTextSizeH=1.000000000000
ModuleTextSizeThickness=0.150000000000
SolderMaskClearance=0.000000000000
SolderMaskMinWidth=0.000000000000
DrawSegmentWidth=0.200000000000
BoardOutlineThickness=0.100000000000
ModuleOutlineThickness=0.150000000000
[cvpcb]
version=1
NetIExt=net
[eeschema]
version=1
LibDir=
[eeschema/libraries]
[schematic_editor]
version=1
PageLayoutDescrFile=
PlotDirectoryName=./
SubpartIdSeparator=0
SubpartFirstId=65
NetFmtName=
SpiceAjustPassiveValues=0
LabSize=50
ERC_TestSimilarLabels=1
```
